# ERASE: a novel surface reconditioning strategy for single-molecule experiments

**DOI:** 10.1101/282632

**Authors:** D W Bo Broadwater, Roger B Altman, Scott C Blanchard, Harold D Kim

## Abstract

While surface-based single-molecule experiments have revolutionized our understanding of biology and biomolecules, the workflow in preparing for such experiments, especially surface cleaning and functionalization remains labor-intensive and time-consuming. Even worse, meticulously assembled flow channels can be used only once for most experiments. A reusable surface would thus dramatically increase productivity and efficiency of single-molecule experiments. In this paper, we report a novel surface reconditioning strategy termed ERASE (Epitaxial Removal Aided by Strand Exchange) that allows a single flow cell to be used for vast repetition of single-molecule experiments. In this method, biomolecules immobilized to the surface through a nucleic acid duplex are liberated when a competing DNA strand disrupts the duplex via toehold-mediated strand displacement. We demonstrate the wide-range applicability of this method with various common surface preparation techniques, fluorescent dyes, and biomolecules including the bacterial ribosome. Beyond time and cost savings, we also show ERASE can assort molecules based on a nucleic acid barcode sequence, thus allowing experiments on different molecules in parallel. Our method increases the utility of prepared surfaces and is a significant improvement to the current single-use paradigm.

## Main

Surface-based, single-molecule experiments comprise an essential toolkit to study biophysical mechanisms. In these experiments, the molecules of interest are tethered to a glass surface via a strong cohesive interaction between the surface and the molecule. Complexes that contain a nucleic acid component are often tethered via hybridization of the nucleic acid component to a single-stranded DNA bound to the surface. Such a tethering scheme has been used for many important biological systems, which include nucleosomes (1-2), polymerases (3-4), endonucleases (5), ribosomes (6-10), ribozymes (11), riboswitches (12), the DNA mismatch repair system (13-15), and CRISPR/Cas9 (16-17).

The power of single-molecule investigation is best harnessed when molecules are housed inside a flow chamber rigged with an inlet and an outlet for liquid delivery. One of the advantages of this setup is the capability to change the solution phase by simple perfusion. Reversible reaction rates can then be measured from surface-bound molecules in various buffer conditions using the same flow channel. However, irreversible reaction steps such as high-affinity ligand binding and ATP or GTP hydrolysis can only be measured once from the molecules within the field of view, which constitutes only a tiny fraction of the flow chamber surface. In this regard, the flow chamber is heavily underutilized.

Similarly, host complexes of a different composition or sequence must often be investigated in separate flow channels (17-18) unless they are modified with a different fluorescent label. Therefore, researchers have to prepare multiple flow channels to experiment with variants of the host complex or to obtain statistics for rate measurements. In our experience, preparing flow channels remains one of the most labor-intensive steps of the experimental procedure. A simple wash protocol that allows repeated usage of a single flow channel for different molecules would thus be highly beneficial to the single-molecule biophysics community.

In pursuit of such strategy, we noted that even a stable DNA duplex can be completely disrupted by a reaction called toehold-mediated strand displacement (18). In this reaction, a single-stranded DNA molecule called the “invader” strand hybridizes to the substrate DNA strand in the toehold region and unzips the adjacent incumbent DNA strand by way of branch migration. We reasoned that this switch functionality could be harnessed as a surface reformatting tool, which would reduce the amount of time and materials necessary for preparing new flow cells.

Here, we introduce a novel surface clearing strategy termed ERASE (Epitaxial Reformatting Aided by Strand Exchange). This method (Fig. 1a) involves three nucleic acid oligomers referred to as anchor, tether, and switch. The anchor is a short (10-15 nt) biotinylated oligomer that serves as the surface attachment site. The tether is the extension of the nucleic acid component of the complex of interest, which is purposefully designed to contain a terminal domain complementary to the anchor followed by a short (7-15 nt) spacer domain. The switch is fully complementary to the spacer and anchor domain of the tether molecule. According to this scheme, the experimental complex is immobilized through base pairing between the anchor and tether, while the addition of the switch removes the surface-tethered complexes through toehold-mediated strand displacement. After displacement, the surface is ready for another experiment by introducing molecules with the tether.

**Figure 1:**
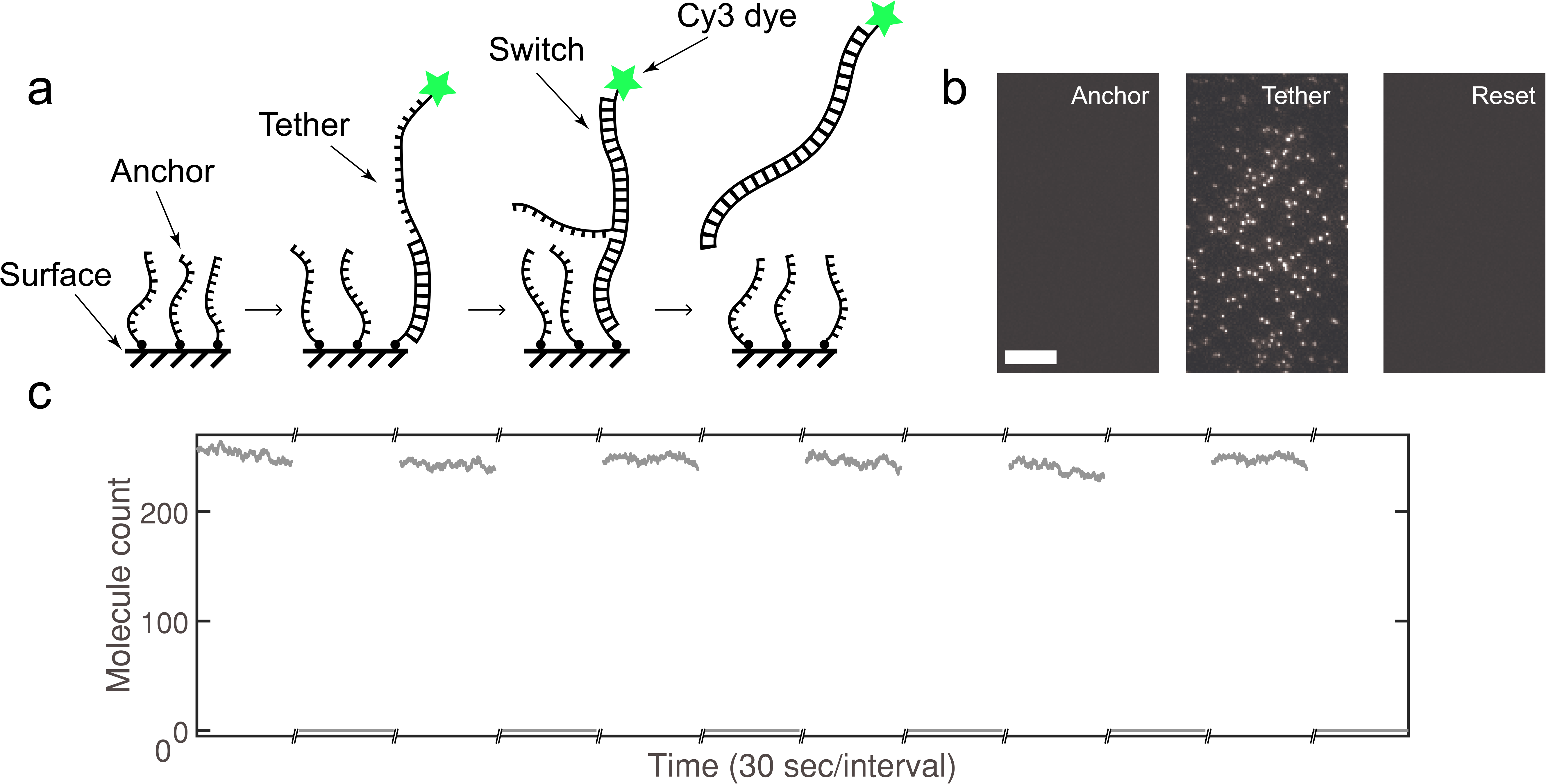
ERASE in action. (a) Biotinylated anchor molecules are immobilized on the surface. Cy3 labeled molecules with a tether are pumped into the flow cell and hybridize with the anchors. After reaching steady-state, switch molecules are flowed in. Strand displacement begins by binding of the switch to the open toehold region of the tether molecule. After branch migration, the anchor is displaced and the tether-switch duplex returns to solution. (b) After annealing to the anchor molecules, the tether molecules reach a steady-state surface density. The switch molecules entirely remove the anchor-bound tether molecules allowing the experiment to be repeated. The scale bar is 10 μm. (c) Molecule counts over 30 second intervals from the same field of view are plotted. The high molecule count is maintained across several rounds of ERASE while the switch molecule efficiently removes all tether molecules.

To demonstrate this method, we first tried a Cy3 labeled single-stranded DNA oligonucleotide containing a tether and a shorter biotinylated DNA strand as the anchor (Fig. 1a). The anchor molecules were introduced at ∼1 nM, which would correspond to a density of 1 anchor/(130 nm)^2^ at full adsorption. This surface density is high enough to provide ample binding sites for the tether, but low enough to prevent any nearest-neighbor interaction. The tether was then introduced at 50 pM, which resulted in an increasing number of spots in the field of view. The time-dependence of this increase could be fit to a negative single exponential model (Supplementary Fig. 1), which suggests that tethers bind to the surface via densely populated anchors in a pseudo first order process.

We next performed ERASE on the flow channel by introducing the switch at 500 nM. The number of spots quickly decayed to the background level over a few seconds (Fig. 1b, Supplementary Fig. 3). As shown in Fig. 1c, the number of spots in the same field of view changed from ∼250 to nearly zero, indicating that ERASE completely removed all bound tether molecules. After ERASE, re-introduction of tethers at 50 pM could regenerate a similar density of spots in the same field of view. We performed this cycle of ERASE and tether addition as many as 6 times, and for each cycle, were able to achieve complete removal and recovery of spots in the same field of view with no sign of fatigue. The rates associated with removal and recovery also showed little difference between all trials (Supplementary Fig. 2). Finally, we found ERASE to be highly sequence-specific as the surface was reformatted only when the complementary sequence was used (Supplementary Fig. 3).

**Figure 2:**
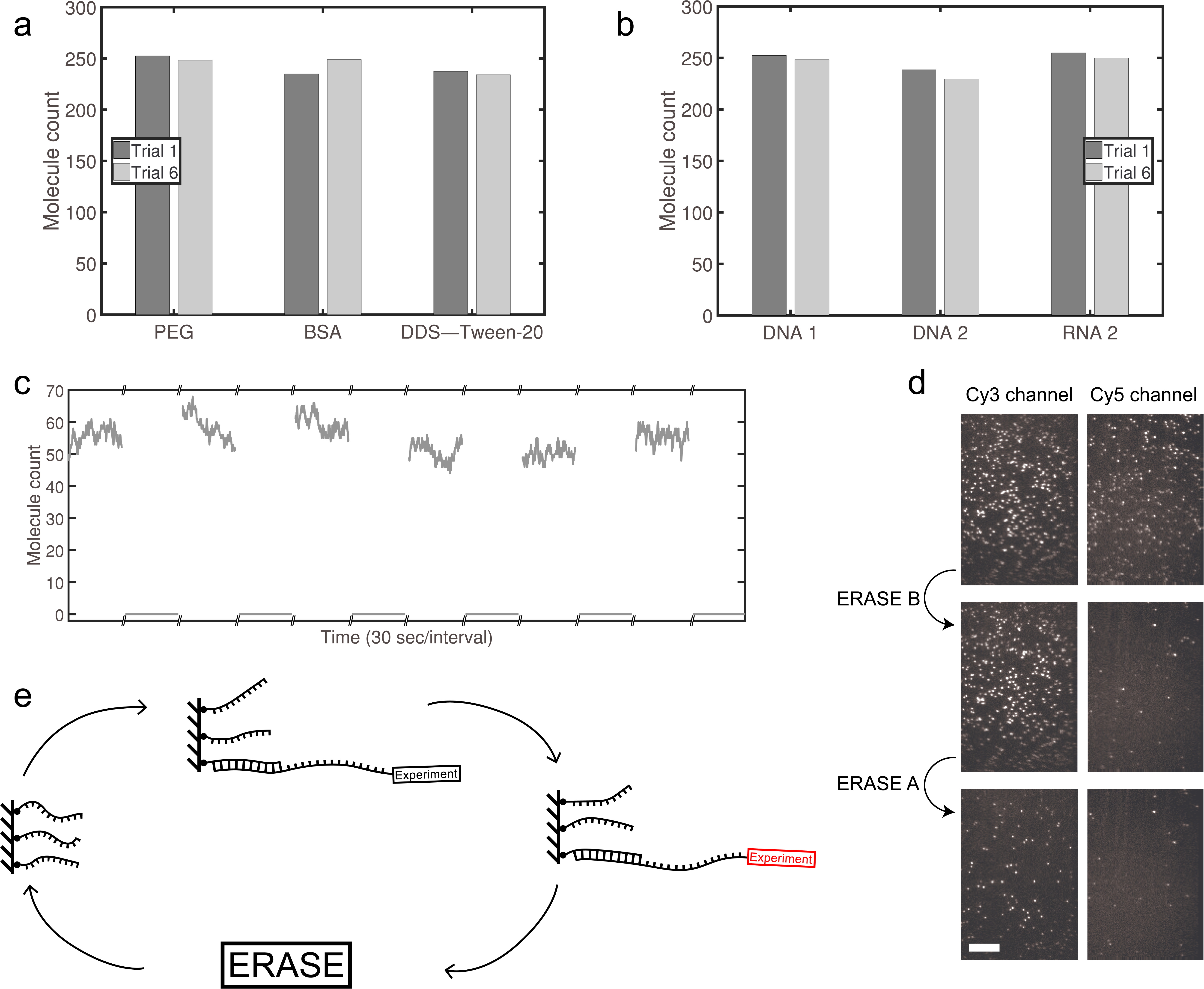
ERASE is versatile. (a) Molecule counts are repeatable and consistent across 3 different surface passivation techniques: PEGylation, BSA, and DDS–Tween-20+BSA. (b) ERASE is highly modifiable and can be constructed with any sequence demanded by the experiment. The molecule counts of a different anchor-tether-switch scheme perform comparably to the first scheme. ERASE is adaptable and can be augmented to different types of nucleic acid. Although it is composed of RNA, the tether molecules can still be entirely removed by the same DNA switch molecules. (c) Molecule counts of 70S ribosome complexes with an RNA tether are plotted, and ERASE efficiently removes all molecules. Molecule counts over the same field of view are consistent across several rounds of ERASE. (d) We designed two different DNA tethers, *A* and *B*: *A* is labeled with Cy3 dye and barcode sequence *a*, and *B* with Cy5 dye and barcode sequence *b*. Both tethers contain a common sequence complementary to the anchor. The Cy3- and Cy5-channel images of the same field of view are shown on the left and right columns, respectively. The first row of images are obtained after incubating the anchor-coated surface with a 1:1 mixture of *A* and *B*, the second row after ERASE with switch *Ā*, which is complementary to *A*, and the third row after ERASE with switch 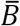, which is complementary to *B*. To minimize signal loss from the corresponding channel, we introduced switch molecules at a lower concentration which decreased ERASE efficiency. The scale bar is 10 μm. (e) ERASE is generalizable to any experiment using a nucleic acid tether. The tether-containing molecule can undergo irreversible changes (red) such as protein modification or annealing; however, provided the toehold remains intact, the switch molecule can always remove the tether-containing molecule.

To showcase the wide adaptability of ERASE, we repeated the experiment with three common surface passivation techniques: PEGylation (19), bovine serum albumin (BSA) (18), and DDS– Tween-20+BSA (20). We observed repeatable performance between trials that was consistent across all passivation schemes (Fig. 2a, Supplementary Fig. 4). ERASE is also adaptable to modifications in sequence as well as anchor and toehold length. We repeated the hybridization experiment with a separately designed scheme containing a different anchor length, toehold length, and sequence. We found the new Cy3 labeled DNA tether to produce a similar steady-state molecule count (Fig. 2b) as well as similar binding kinetics (Supplemental Fig. 5). Again, the switch molecule completely removed tethers from the surface. Finally, ERASE can be expanded to other nucleic acid types. We performed the experiment with a Cy5-labeled RNA tether with the same sequence as the second DNA tether except T-to-U substitution and observed comparable performance (Fig. 2b, Supplemental Fig. 5). Despite being a different nucleic acid, the same DNA switch molecule from the second design scheme efficiently removed the RNA tether.

The true power of ERASE lies in the ability to reprogram the surface for any biomolecules that harbor an end-exposed nucleic acid tether. To demonstrate this, we repeatedly ERASEd 70S ribosome particles (Cy3-Met-tRNA^fMet^ in the P site) containing an mRNA tether. ERASE consistently removed all ribosomal complexes while molecule counts for each round were very similar (Fig. 2c). The switch molecule, designed to hybridize the mRNA just beyond the anti Shine-Dalgarno sequence (Supplementary Table 1), specifically removed ribosomal complexes whereas a bioorthogonal control switch did not (Supplementary Fig. 6).

Finally, we explored one possible application of ERASE for multiplexing. In this scheme, multiple types of molecules carrying different barcodes in the toehold region would be immobilized to the same flow channel and subjected to the same perfusion experiment. Subsequently, the positions of each molecule type could be determined by performing ERASE with barcode-specific switches in a sequential manner, and the mixed signal be decomposed. As a proof of principle, we immobilized two different tether molecules fluorescently labeled with Cy3 and Cy5, respectively, and performed ERASE with one switch at a time. Each switch removed ∼95% of its target molecules while non-target molecules were removed at a much lower rate of 1% (Fig. 2d). Optimizing the ERASE condition to maximize the false negative rate and minimize the false positive rate will be the topic of a future study.

Apart from the time and cost savings due to repeated use of flow cells, ERASE provides several practical advantages to the experimenter. In case inhomogeneities from channel to channel or from field to field become an issue, ERASE allows the same field of view to be reused across all experiments. Further, since switch molecules specifically disrupt base pairing between the anchor and the tether, spots that are not removed by switch molecules can be identified as nonspecifically bound or spurious and, hence, can be left out of data processing and statistics (Fig. 2d). Therefore, ERASE provides an *a posteriori* mechanism to select “good” molecules in an unbiased manner.

In conclusion, we introduced ERASE, a novel surface reformatting scheme that enhances the power of surface-based, single-molecule experiments. We have shown ERASE to be highly specific for removing the target molecules from the surface without compromising the long-term binding capacity of the surface for subsequent experiments. We have also shown it to be consistent and repeatable across common surface passivation techniques, and different sequences and types of nucleic acid. Due to its simplicity and flexibility, we expect ERASE to become routine for future single-molecule studies.

## Methods

### Sample Preparation

Custom nucleic acid oligomers were ordered from Integrated DNA Technologies and modified to include a Cy3 or Cy5 fluorophore near the 5’ end or a biotin linker at either the 5’ or 3’ end. Unmodified switch molecules were ordered from Eurofins Scientific. The specific sequences are in Supplementary Table 1. 70S ribosomal complexes were prepared as described previously (7).

### Slide and Coverslip Cleaning and Functionalization

8 holes were drilled by computer numerical control (CNC) across both long edges of a quartz slide using a diamond drill bit. Quartz slides and coverslips were sonicated for 10 minutes in Milli-Q water and dried by vacuum for 10 minutes. Slides and coverslips were then placed in an upright position and plasma cleaned for 10 minutes (Harrick Plasma, PDC-32G). Flow cells were constructed by laying thin strips of double-stick tape across the slide. After aligning and pressing the coverslip against the slide, the open edges were sealed with 5-minute epoxy. Three surface functionalization methods were performed: 1) BSA, 2) PEGylation, and 3) DDS–Tween-20+BSA. 1) After slide construction, 25 μL biotinylated BSA (1 mg/mL) was incubated for 5 minutes in the flow cell. 2) Prior to flow cell construction, 2 mg of biotin-PEG-silane was mixed with 80 mg of mPEG-silane into 320 μL of 100 mM sodium bicarbonate solution. 80 μL of PEG solution was poured onto the quartz slide. The coverslip was placed on top of the slide and both were incubated at room temperature for an hour. The pair was disassembled, washed with Milli-Q water, and dried with compressed air. Flow cells were then assembled as described above. 3) After construction, slides and coverslips were rinsed in hexane and incubated in an upright container with a solution containing 75 mL hexane and 50 μL dichlorodimethylsilane for 1.5 hours while gently shaking. The slides and coverslips were sonicated 3 times in fresh hexane for 2 minutes each. After slide construction, 25 μL biotinylated BSA (1 mg/mL) was incubated for 5 minutes in the flow cell. Then 100 μL 0.2% Tween-20 in T50 (10 mM Tris-HCl, 50 mM NaCl, pH 8.0) was incubated for 10 minutes in the flow cell. After all 3 passivation techniques, the surface was neutravidin coated.

### Experimental Setup

Objective-type total internal reflection fluorescence microscopy was performed to image single molecules with a commercial microscope (Olympus IX81). Cy3 labeled molecules were excited by 532-nm laser (NT66-968, B&W Tek), and Cy5 labeled molecules were excited by 640-nm laser (CUBE 640-30FP, Coherent). Images were captured by EMCCD (DU-897ECS0-#BV, Andor Technology) at 100 ms exposure and 2×2 binned by our in-house software. A syringe pump was (NE-1000, New Era Pump System) used to control flow volume and flow rate (15 μL/s). Nucleic acid molecules were pumped into flow cells in oxygen-scavenging imaging buffer containing 1 mM 6-hydroxy-2,5,7,8-tetramethylchroman-2-carboxylic acid (Trolox), 5 mM protocatechuic acid, 100 nM protocatechuate 3,4-dioxygenase, 100 mM Tris-HCl (pH 7), and 1 M NaCl. Experiments with 70S ribosome complexes were performed in Tris-polymix buffer containing 50 mM Tris-acetate (pH 7.5), 5 mM MgCl_2_, 100 mM KCl, 5 mM NH4(CH_3_COO),0.5 mM CaCl_2_, 0.1 mM EDTA, 5 mM putrescine, and 1 mM spermidine in the presence of an oxygen-scavenging buffer consisting of 2 mM protocatechuic acid, 50 nM protocatechuate 3,4-dioxygenase, and 1 mM Trolox. The fluorescent signal was recorded and analyzed using in-house MATLAB software. Molecules were counted by peak detection.

### Contributions

D.W.B.B. and H.D.K. conceived and designed experiments. D.W.B.B. performed the experiments and analyzed the data. R.B.A. and S.C.B. constructed the ribosome complex. All authors contributed in writing the manuscript.

