## Supplementary Materials for "ERASE: a novel surface reconditioning strategy for single-molecule experiments"

### Supplementary information

| | |
|---|---|
| **Anchor 1** | /BioTEG/CACTGGTGTT |
| **Tether 1** | CA/iCy3/ATTAAAATTCCGACAACACCAGGT |
| **Switch 1** | ACCTGGTGTTGTCGGAATTTTAAT |
| **Control Switch 1** | GAGTGGGGAGTCAAAGTAAATTCAAAACCAGGACTCACTGCGAGGTA |
| **Anchor 2** | TCAATTCGTCGTC/BioTEG/ |
| **Tether 2** | /Cy3/GACGACGAATTGAAGTGAAA |
| **Switch 2** | TTTCACTTCAATTCGTCGTC |
| **RNA Tether 2** | /Cy5/GACGACGAAUUGAAGUGAAA |
| **DNA Tether-70S** | AGTTTTAGGTTGCCCCCCTTTTTTTTTTTTTTTTTTTTTTTTTTTTTT/BioTEG/ |
| **mRNA-70S** | CAACCUAAAACUUACACACCCUUAGAGGGACAAUCGAUGUUCAAAGUCUUCAAAGUCAUC |
| **Switch-70S** | GGGTGTGTAAGTTTTAGGTTG |
| **Orthogonal Tether** | /CY5/GACGACGAATTGATCACTTT |
| **Orthogonal Switch** | AAAGTGATCAATTCGTCGTC |

**Supplementary Table 1**

**Oligonucleotide sequences used in this study**

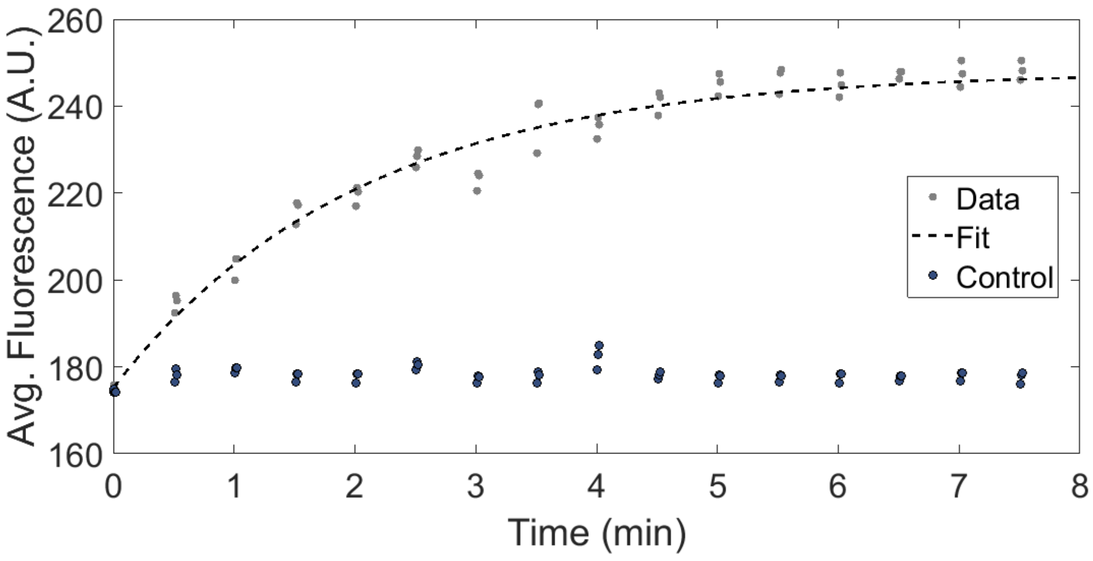

**Supplementary Figure 1**

**Mean fluorescence from Cy3 labeled DNA binding to a PEGylated surface**

To avoid photobleaching the laser is strobed for 2 second intervals every 30 seconds at an exposure time of 500 ms. The data is fitted with a single exponential to obtain a rate of anchor hybridization. As a control, Cy3 labeled DNA is introduced to a surface without anchor molecules.

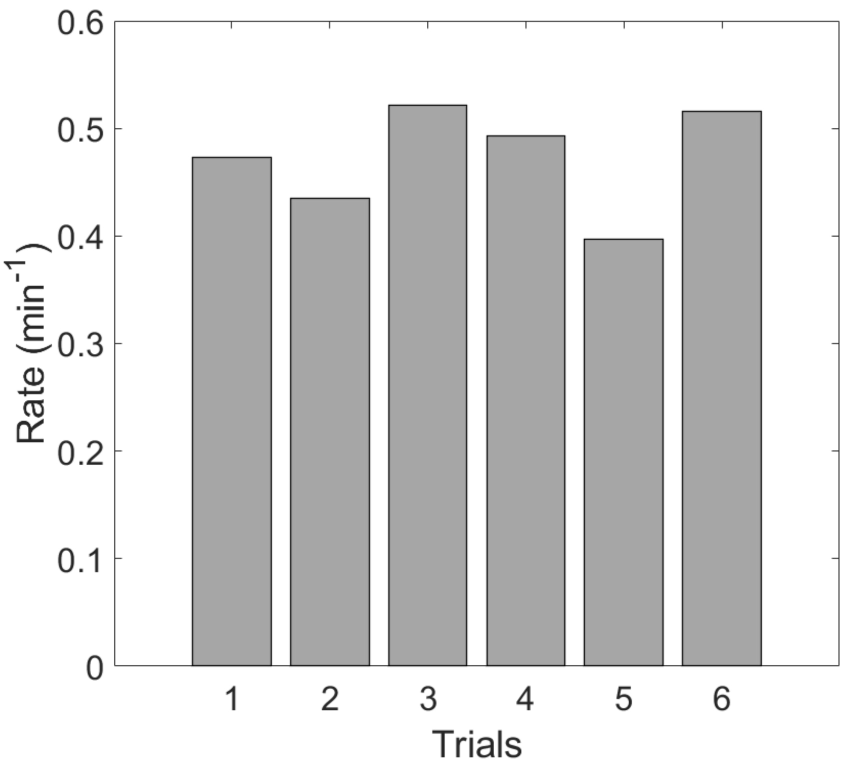


**Supplementary Figure 2**

**Rate of mean fluorescence over rounds of ERASE**

Similar rate constants were extracted from single exponential fits of anchor hybridization for several consecutive rounds of ERASE.

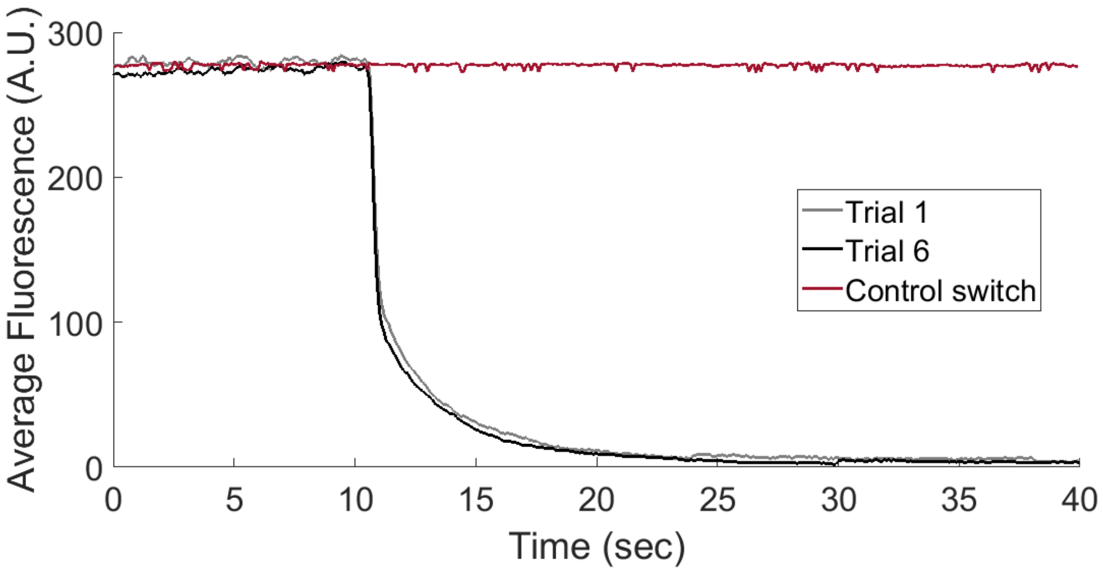


**Supplementary Figure 3**

**Rate of mean fluorescence for Cy3 labeled DNA after introducing switch**

The switch molecule is introduced at 10 seconds. Decrease in fluorescence is complete in tens of seconds and is similar from Trial 1 to Trial 6. A control switch with a different sequence does not engender a change in fluorescence.

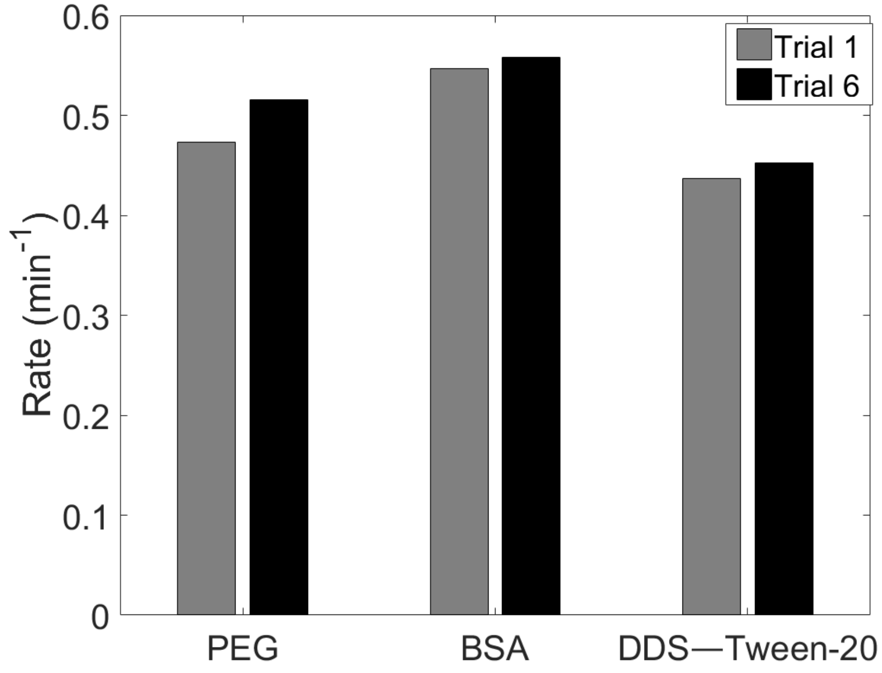


**Supplementary Figure 4**

**Rate of mean fluorescence for Cy3 labeled DNA across different surface passivation schemes**

Similar rate constants were extracted from single exponential fits of anchor hybridization for the first and sixth round of ERASE for three surface passivation techniques: 1) PEGylation, 2) BSA, 3) DDS–Tween-20+BSA.

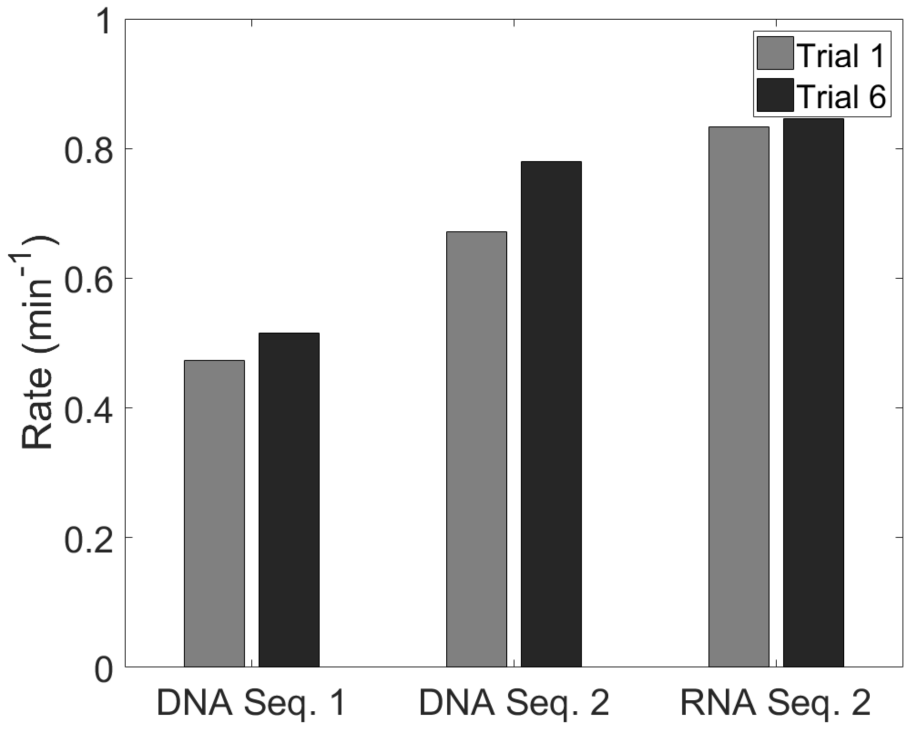


**Supplementary Figure 5**

**Rate of mean fluorescence different sequences of Cy3 labeled DNA and Cy5 labeled RNA**

Similar rate constants were extracted from single exponential fits of anchor hybridization for the first and sixth round of ERASE for different nucleic acids: 1) DNA with internally labeled Cy3, 2) DNA with a different sequence and end labeled with Cy3, 3) RNA with the same sequence as 2) and end labeled with Cy5.

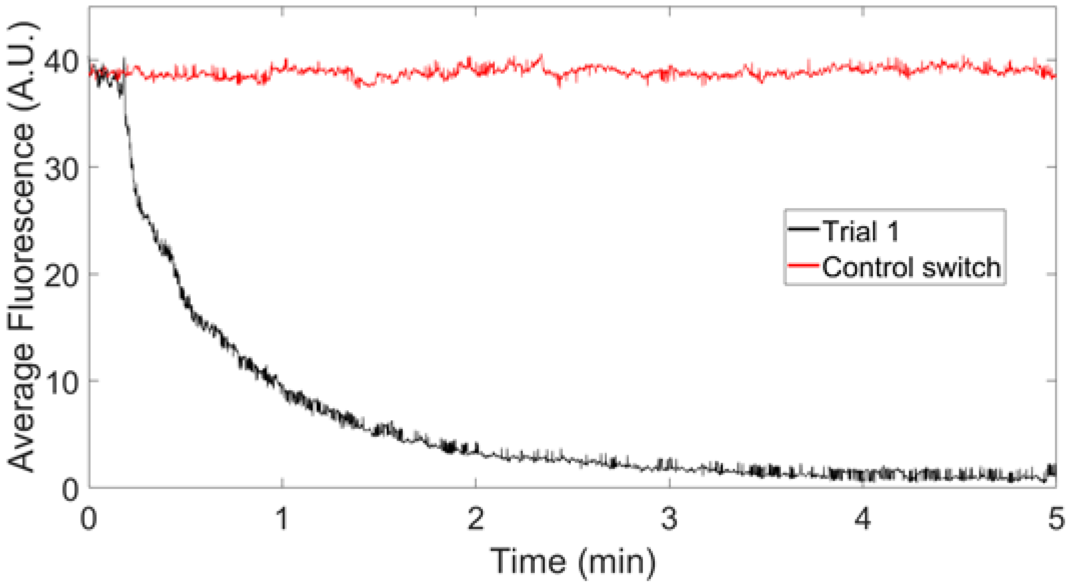


**Supplementary Figure 6**

**Rate of mean fluorescence for 70S complexes after introducing switch**

The switch molecule is introduced at 10 seconds. Decrease in fluorescence is complete in a few minutes. A control switch with a different sequence does not cause a change in fluorescence.
